# Draft genome of the aquatic moss *Fontinalis antipyretica* (Fontinalaceae, Bryophyta)

**DOI:** 10.1101/2020.04.29.069583

**Authors:** Jin Yu, Linzhou Li, Sibo Wang, Shanshan Dong, Ziqiang Chen, Nikisha Patel, Bernard Goffinet, Huan Liu, Yang Liu

## Abstract

**Background:** Mosses compose one of the three lineages that form the sister group to extant vascular plants. Having emerged from an early split in the diversification of embryophytes, mosses may offer complementary insights into the evolution of traits following the transition to and colonization of land. Here, we report the draft nuclear genome of *Fontinalis antipyretica* (Fontinalaceae, Hypnales), a charismatic aquatic moss widespread in temperate regions of the Northern Hemisphere. We sequenced and *de novo* assembled its genome using the 10 × genomics method. The genome comprises 486.3 Mb, with a scaffold N50 of 38.8 kb. The assembly captured 89.4% of the 303 genes in the BUSCO eukaryote dataset. The newly generated *F. antipyretica* genome is the third genome of mosses, and the second genome for a seedless aquatic plant.

## Introduction

With ~13,000 extant species, mosses represent perhaps the second most speciose lineage of land plants [1]. Mosses diverged from their common ancestor with liverworts (One Thousand Plant Transcriptomes Initiative, 2019) no later than 350 Mya [2–4]. The early diversification of land plants is marked by various morphological innovations, such as branching of the sporophyte or stomata [5], as well as metabolic innovations, most notably perhaps biopolymers, essential for composing the cuticle [6] that enables plants adapt to a living environment featured by water-deficiency and UV-exposure. To date, two nuclear genomes have been sequenced for mosses, namely the model taxon and acrocarpous moss *Physcomitrium patens* [7], and *Pleurozium schreberi* [8], a representative of the diverse pleurocarpous hypnalean mosses.

*Fontinalis antipyretica* (NCBI: txid67435) is an aquatic moss species (Fig. 1) from the most diverse moss order, i.e., the Hypnales [9]. Sequencing the genome of *F. antipyretica* should provide the first opportunity for a comparative genomic study in this lineage that may have diversified after the rise of the angiosperms. Furthermore, this is the second genome for a seedless aquatic plant, it will also allow for the assessment of independent genomic transformations linked to a reversed shift to an aquatic habitat. Thus the genome of this species would contribute to the framework necessary to study genome evolution in mosses, and to explore the adaptive transformations underlying the shifts between terrestrial and aquatic habitats.

**Figure 1.**
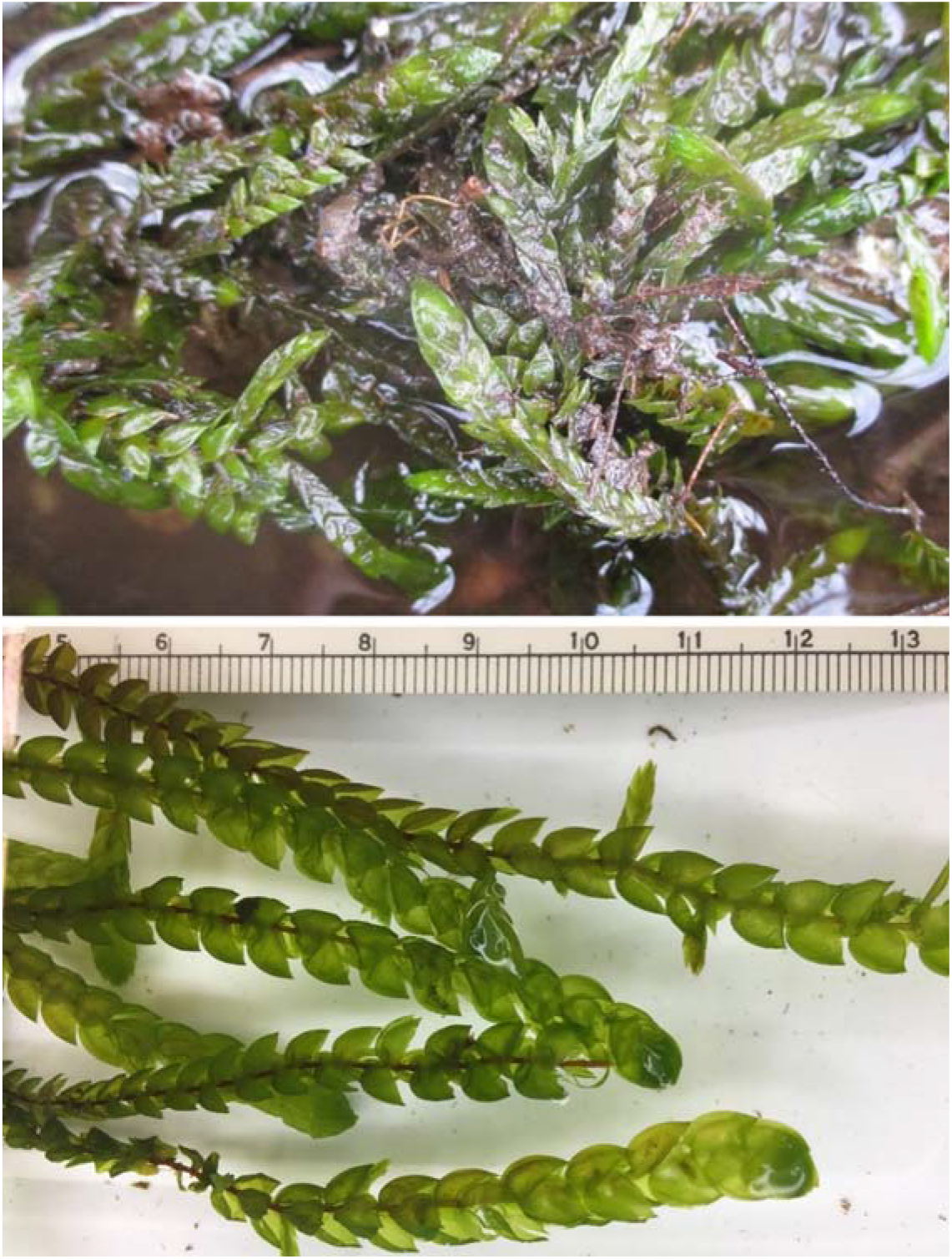
Photographs of the aquatic moss Fontinalis antipyretica. Upper: a wild population; lower: shoots with a scale (in cm).

## Methods

Fresh gametophyte tissue of *Fontinalis antipyretica* was collected in Connecticut, USA, and the voucher specimen (Collection number: Goffinet 14197) has been deposited in the George Safford Torrey Herbarium at the University of Connecticut (CONN). The genomic DNA was extracted at the Fairy Lake Botanical Garden, and has been deposited with the DNA extraction number of 332.

The plant tissue was cleaned under a dissecting microscope to enhance the quality of the material. Approximately 0.4 g fresh plant shoots were ground in liquid nitrogen, and used for DNA extraction using the NucleoSpin Plant midi DNA extraction kit following the manufacture’s protocol (Macherey-Nagel, Düren, Germany). The genomic DNA was quality controlled using a Qubit® 3.0 Fluorometer (Thermo Fisher Scientific, USA). The high molecular weight genomic DNA was used to construct the 10× Genomics libraries [10] with insert sizes of 350–500 bp following the manufacturer’s protocol (Chromium Genome Chip Kit v1, PN - 120229, 10× Genomics, Pleasanton, USA) [11]. The libraries were sequenced on a BGISEQ-500 sequencer (RRID:SCR_017979) to generate the 150 bp paired-end reads [12]. The raw reads were directly used for genome assembly using 10× Genomics Supernova v2.1.1 (RRID:SCR_016756) with auto filtering of the reads [10].

For the genome assembly, we first calculated the distribution frequency of the barcodes in the raw data, and removed those reads containing barcodes with extremely low or high frequencies. The remaining reads were subsequently de novo assembled using 10× Genomics Supernova. Then, we used GapCloser v1.12-r6 (RRID: SCR_015026) to close the gaps of the preliminary assembly [13]. All these softwares were used with default parameters.

The genome size of *Fontinalis antipyretica* was estimated using flow cytometry. Mature leaf tissue of *Raphanus sativus* L. cv. Saxa was used for internal and external standardization. *R. sativus* has an established 2C genome size of 1.11 pg [14] and was cultivated from seeds obtained from the Institute of Experimental Botany (Olomouc, Czech Republic). Two assays were externally standardized and one assay was internally standardized. For each, 0.2 g of fresh tissue from the sample or the standard were used. Fresh tissue was combined with 750 μl of Cystain PI Absolute P nuclei extraction Buffer (Sysmex, Kobe, Japan) in a glass petri dish maintained on ice and chopped with a clean razor blade for 60 seconds. The internally standardized sample was co-chopped with tissue of the standard, *R. sativus*. The resulting nuclear suspension was transferred to a 30 μm CellTrics filter (Sysmex, Kobe, Japan). The flow through was combined with 500 μl of Cystain PI Absolute P staining solution (Sysmex, Kobe, Japan), 150 μg/mL of propidium iodide, and 50 μg/mL of RNAse. Samples were incubated on ice for 30–60 minutes. Flow cytometry was run on a BD Biosciences LSRFortessa X-20 Cell Analyzer.

The cytometry data were visualized using FlowJo™ v10.6.2 software (FlowJo, LLC, Ashland, OR, USA). For each assay, to estimate genome size, the 1C nuclei of *F. antipyretica* were compared to the 2C nuclei of *Raphanus sativus*. The ratio of the mean fluorescence of the 1C F. antipyretica peak and the *R. sativus* 2C peak was multiplied by the genome size of *R. sativus*. The genome size estimate produced here is the mean of the estimates produced by the two externally standardized assays as well as the one internally standardized assay.

For genome annotation, we used Piler v1.0 (RRID:SCR_017333) [15], Repeatscout v1.0.5 (RRID:SCR_014653) [16], LTR Finder v1.0.6 (RRID:SCR_015247) [17] and RepeatMasker v4.0.6 (RRID:SCR_012954) [18] to conduct *de novo* repeat element prediction. RepeatMasker v4.0.6 was also implemented for identification of repeats based on known repetitive sequences. Gene structure annotation was performed by the MAKER v2.31.8 (RRID:SCR_005309) [19] pipeline, integrating results from ab initio gene predictors, EST evidence, and protein homologs. Augustus v3.2.1 (RRID:SCR_015981) [20], GeneMark v4.32 (RRID:SCR_011930) [21] and SNAP v2006-07-28 (RRID:005501) [22] were used for *ab initio* gene prediction. Transcriptome assembly of *F. antipyretica* was obtained from the one-kp project and used as EST evidence. Protein sequences from closely-related green plants were selected as homolog-based evidence.

To reconstruct the phylogenetic tree, we used OrthoFinderv2.3.7 (RRID:SCR_017118) to search for single-copy orthologs among the genomes of *F. antipyretica* and eight other green plants, including *Klebsormidium nitens*, *Chara braunii*, *Anthoceros angustus*, *Marchantia polymorpha*, *Sphagnum fallax*, *Physcomitrium patens*, *Pleurozium schreberi*, and *Selaginella moellendorffii*. The genomes were downloaded from Phytozome database (https://phytozome.jgi.doe.gov/). A total of 472 single-copy loci were selected, and each locus was aligned by MAFFT v7.3.10 (RRID:SCR_011811) [25], and concatenated into one super-matrix. Finally, RAxML v8.2.4 (RRID:SCR_006086) was implemented to construct the maximum likelihood tree, using the PROTCATGTR substitution model [26]. The result tree was uploaded to iTOL (https://itol.embl.de/) for visualization.

## Results and Discussion

### Genome assembly and annotation

A total of 133 Gb PE150 raw data were generated by the BGISEQ-500 sequencer. The genome of *F. antipyretica* totaled 486.3 Mb spanning 154,086 contigs, with a contig N50 of 24.6 kb. The final scaffold assembly included 130,576 scaffolds with a N50 length of 38.8 kb. Our assembly captured 89.4% of the 303 genes in the BUSCO eukaryota dataset [27].

The GC content of *F. antipyretica* is 43.91%, which is higher than that of *Physcomitrium patens* (i.e., 33% [7]), or *Pleurozium schreberi* (26.4% [8]). The size of the genome of *F. antipyretica* is estimated at 486.3 Mb, which is comparable to the size of *P. patens*’ genome (i.e., 462.3 Mb), but larger than that of *P. schreberi* (i.e., 318.3 Mb). Repeats make up 42.7% of the *F. antipyretica* genome, compared to 57.0 % in *P. patens* and 28.4% in *P. schreberi*. With 26,398 genes the gene space of the *F. antipyretica* genome is intermediary between *P. patens* with 32,926 genes and *P. schreberi* with 15,992 genes.

### Data validation and quality control

To determine the genome size, we carried out flow cytometry experiment and *k*-mer analysis for *F. antipyretica*. For flow cytometry, the nuclear peaks from which genome size was estimated comprised, on average, 242 events (see Fig. 3 for a representative histogram). The mean coefficient of variance was 7.62. The mean estimated genome size is 0.484 pico-grams. The *k*-mer analysis was performed using the program Jellyfish v2.3.0 (RRID:SCR_005491) with default parameters [28]. The genome size was estimated by dividing the total *k*-mer number by the peak coverage in the *k*-mer distribution curve (Fig. 2). Thus, the genome size was estimated to be ca. 579 Mb, which is slightly larger than the flow cytometry result and genome assembly.

**Figure 2.**
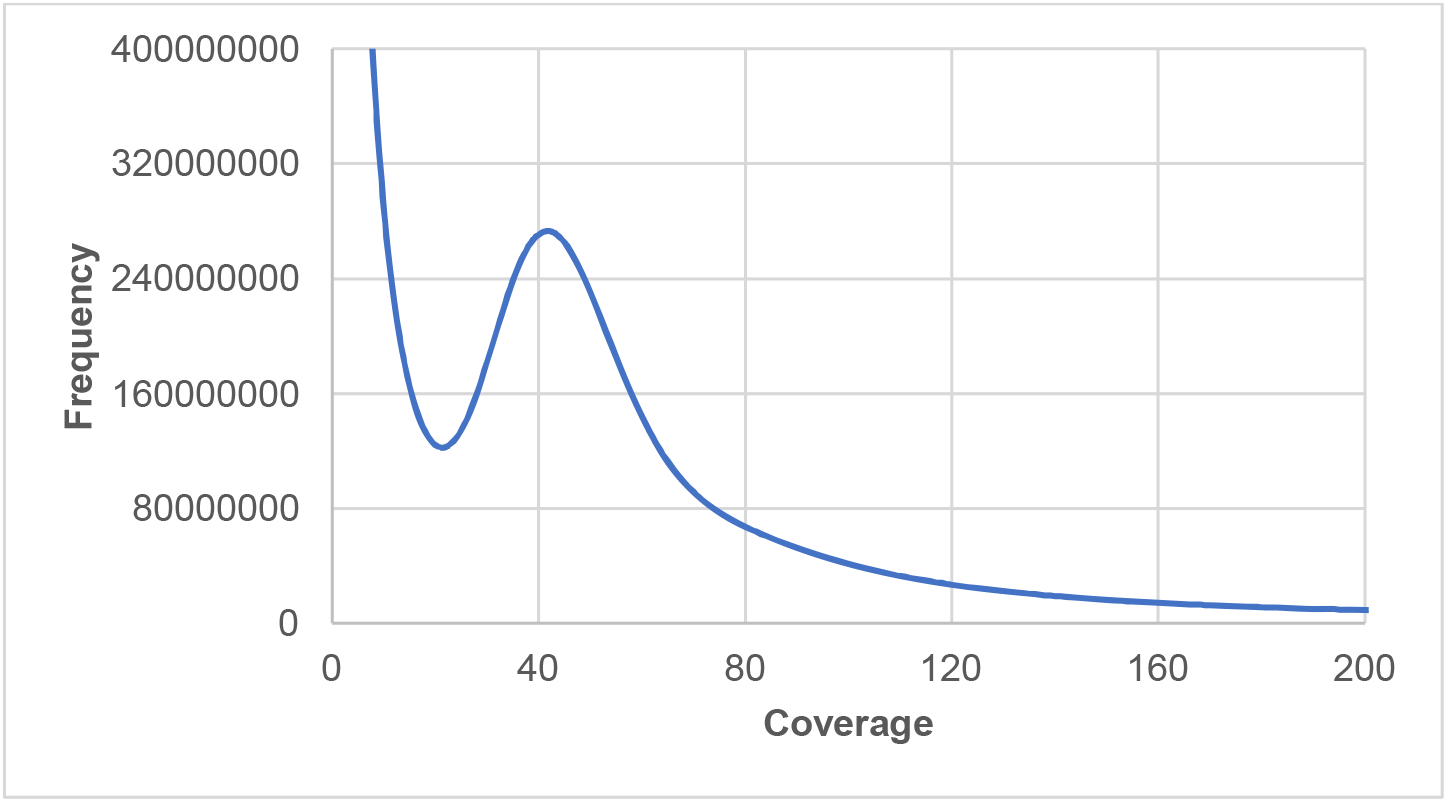
The *k*-mer distribution curve of the *Fontinalis antipyretica* genome data. The curve shows a clear one-peak mode, indicating low heterozygosity and repetitive content across the genome.

**Figure 3.**
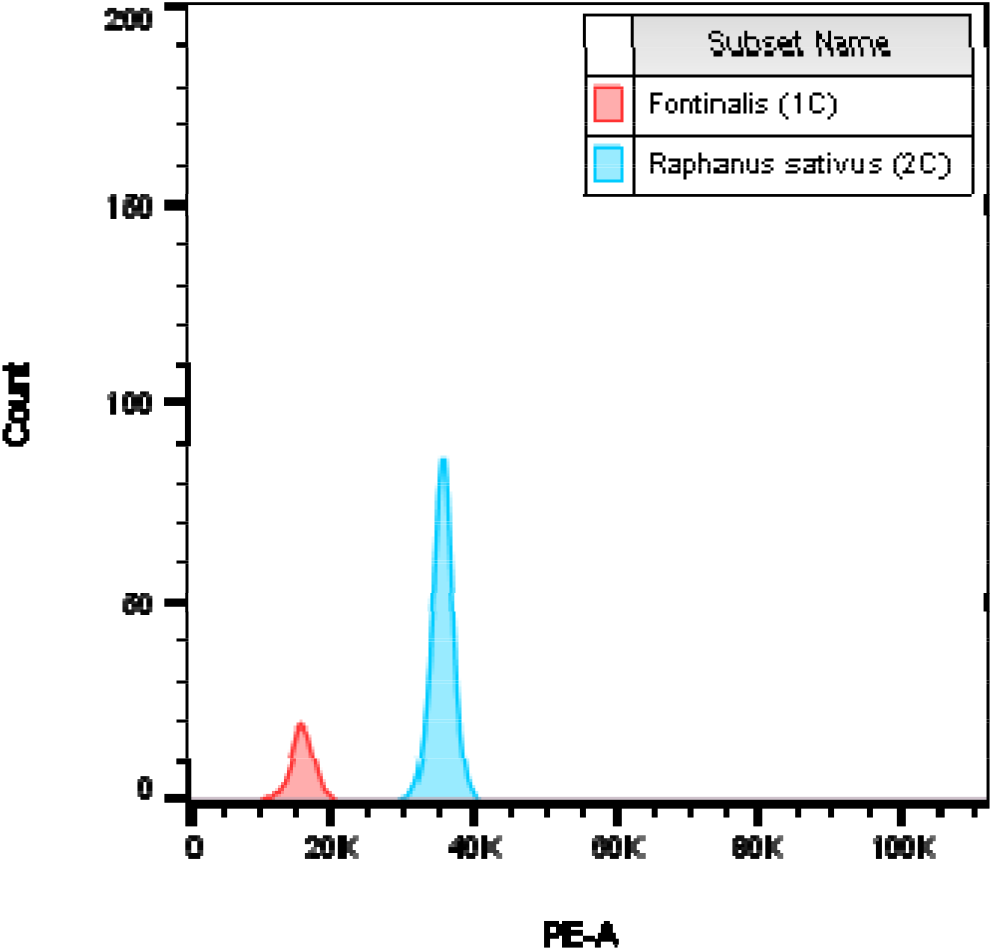
Representative sample of flow cytometry results. The 1C peak of Fontinalis antipyretica and the 2C peak of *Raphanus sativus* cv. Saxa are overlaid to show fluorescent intensity differences on the x-axis indicated by PE-A.

To evaluate the completeness of the assembly, we conducted BUSCO v3.1.0 (RRID:SCR_015008) assessment on the assembly [27]. The assembly captured 89.4% complete BUSCOs of the 303 genes in the BUSCO eukaryote odb9 dataset.

With the streptophyte alga *K. nitens* rooted as the outgroup, bryophytes were confirmed as a monophyletic group, and sister to the vascular plant *S. moellendorffii*. Within bryophytes, hornwort is sister to liverworts and mosses, which is consistent with previous studies [29]. The four mosses formed a clade, the newly sequenced *F. antipyretica* and another Hypnales species *P. schreberi* clustered as one clade (Fig. 4).

**Figure 4.**
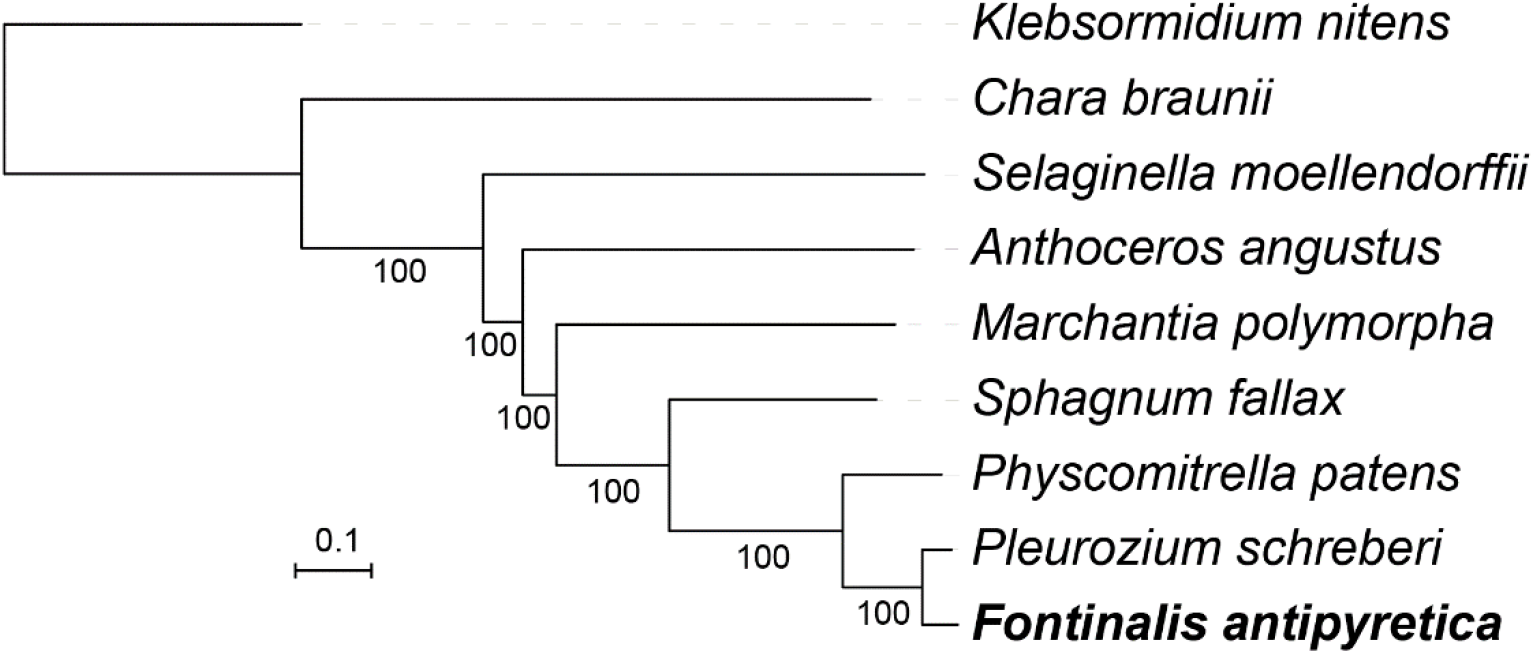
Phylogenetic tree reconstructed using nuclear genome single-copy genes showing phylogenetic relationship of *F. antipyretica* and eight other green plants. Numbers below branches are bootstrap support values. The newly sequenced *F. antipyretica* is in bold.

### Re-use potential

The transition of green plants from freshwater habitats to land catalyzed a major biotic diversification, which lead to major climatic changes on earth. The colonization of land is characterized by the acquisition of many key innovations by plants, such as the development of an embryo, a cuticle, gravitropic detection, and pathogen defense, which were likely crucial for plants to survive in terrestrial environments [30]. The accumulation of genomic data, including the assembly of this moss genome, may contribute to reconstructing the evolution of the developmental networks underlying these innovations.

Reconstructions of the relationships of extant land plant lineages are converging on a scenario wherein bryophytes compose a sister lineage to living vascular plants, with mosses and liverworts sharing a unique common ancestor that arose from a split from the ancestor giving rise to hornworts [31]. Following the recent release of the hornwort genomes [29, 32], gene and gene family evolution among bryophytes can be assessed within a robust phylogenetic framework. With the resolution of the relationships among mosses [33], the accumulation of moss genomes will enable more critical estimates of trends in gene family diversity during the diversification of this lineage of land plants. Furthermore, *Fontinalis* is the first aquatic plant with a gametophyte dominated life cycle, to have its genome assembled and annotated, providing a unique opportunity to evaluate similarities in parallel adaptations in mosses, ferns [34] and angiosperms [35] following shifts to freshwater habitats.

## Acknowledgements

The authors would like to thank Yang Peng and Na Li at the Shenzhen Fairy Lake Botanical Garden for lab assistances.

## Availability of supporting data

The raw reads have been deposited in the NCBI Sequence Read Archive (SRA; accession no. PRJNA627325). The sequence reads and assemblies of the *F. antipyretica* genome have been deposited in the China National GeneBank DataBase (CNGBdb) under the accession number of CNP0000847. Genome assemblies, protein-coding genes, and repeat annotations have been deposited in the GigaScience GigaDB [36].

## Author contributions

YL, HL, and BG conceived and designed the study. BG collected the material. SD and ZC performed the experiments. YJ, ZC, LL, SW, and NP carried out the analyses. YJ drafted the manuscript. YJ, DS, YL, and BG revised the manuscript. All authors have read and approved the final manuscript.

